# From Virtual Screens to Cellular Target Engagement: New Small Molecule Ligands for the Immune Checkpoint LAG-3

**DOI:** 10.1101/2024.08.04.604031

**Authors:** Natalie Fuchs, Laura Calvo-Barreiro, Valerij Talagayev, Szymon Pach, Gerhard Wolber, Moustafa T. Gabr

## Abstract

Herein, we performed a virtual screening study to discover new scaffolds for small molecule-based ligands of the immune checkpoint lymphocyte-activation gene 3 (LAG-3). Molecular dynamics (MD) simulations using the LAG-3 structure revealed two putative binding sites for small molecules: the antibody interface and a lipophilic canyon. A 3D pharmacophore screening resulted in the identification of potential ligands for these binding sites and afforded a library of 25 compounds. We then evaluated the screening hits for LAG-3 binding via microscale thermophoresis (MST) and surface plasmon resonance (SPR). Our biophysical screening identified two binders with *K*_D_ values in the low micromolar range, compounds 3 (antibody interface) and **25** (lipophilic canyon). Furthermore, we investigated the ability of LAG-3 hits to engage LAG-3 on a cellular level using a cellular thermal shift assay (CETSA), where compound **3** emerged as a promising candidate for future development.

---

The lymphocyte-activation gene 3 (LAG-3) immune checkpoint became the latest FDA-approved immune checkpoint target in March 2022,^1^ after anti-cytotoxic T lymphocyte associated protein 4 (CTLA-4), antiprogrammed cell death protein 1 (PD-1), and antiprogrammed death-ligand 1 (PD-L1) monoclonal antibodies (mAbs) were previously developed.^2-7^ This later antiLAG-3 therapy, Relatlimab, was approved for the treatment of patients with unresectable or metastatic melanoma in combination with Nivolumab (anti-PD-1), successfully improving clinical efficacy without increasing toxicity compared to previously approved monotherapy.^1^ Over a dozen more agents that target LAG-3, alone or in combination with other immune checkpoints, such as anti-CTLA4, anti-PD-1, or anti-T cell immunoreceptor with immunoglobulin and ITIM domain (TIGIT), among others, are currently being evaluated clinically for cancer immunotherapy.^8^ The rationale behind targeting multiple immune checkpoints is that different molecules expressed on exhausted immune cells regulate distinct components of the immune response, so targeting various immune mechanisms might restore immune cell function against the tumor more effectively.^9^

While a vast number of mAbs targeting hLAG-3 are being investigated in preclinical and clinical studies, further therapeutics, including small molecules, are just starting to be considered as alternatives. The main advantages of small molecules over mAbs are their oral bioavailability, solid tumor penetration, decreased likelihood of causing adverse immune responses over time, and higher accessibility due to low manufacturing costs.^10^ Importantly, the possibility of optimizing a small molecule’s pharmacokinetic properties: absorption, distribution, metabolism and excretion (ADME); can directly relate to toxicity properties and, in turn, avoid immune-related adverse effects.^10^

Our group recently described a first-in-class small molecule, SA-15-P, that successfully inhibits LAG-3/major histocompatibility complex (MHC) class II and LAG-3/fibrinogen-like protein 1 (FGL-1) interactions.^11^ In that study, we first established a diversified chemical library from different commercial and public libraries that allowed us to perform a focused screening and get an initial hit compound. Afterward, we investigated 25 structural analogs using a structure-activity relationship (SAR) approach, resulting in the identification of the small molecule SA-15-P.^11^ Previously, a comparable strategy aimed at the V-domain Ig suppressor of T cell activation (VISTA) im-mune checkpoint proved successful in identifying and optimizing a lead small molecule that binds to and inhibits VISTA activity.^12^

In 2022, the ectodomain structure of hLAG-3 and its interaction with FGL-1 as well as with a potent antagonist antibody was revealed.^13^ This new description gave novel insights into the understanding of the hLAG-3 immune checkpoint and can be of high importance to effectively design hLAG-3-targeted immunotherapies. Certainly, our team has previously leveraged the crystal structure of another immune checkpoint, the inducible T cell costimulator (ICOS), either independently or alongside its natural ligand, ICOS-L, as a blueprint for developing and validating a virtual screening method.^14, 15^ This approach successfully identified a pioneering small molecule ICOS binder.^14^ Given the encouraging results of the preceding strategy with the ICOS protein, we were keen to investigate whether this approach would yield similar success with the LAG-3 immune checkpoint. Therefore, we used a 3D pharmacophore-based virtual screening approach and then evaluated the screening hits’ binding ability to LAG-3. Additionally, we investigated validated binders in a cellular environment. Our workflow is illustrated in Figure 1.

**Figure 1.**
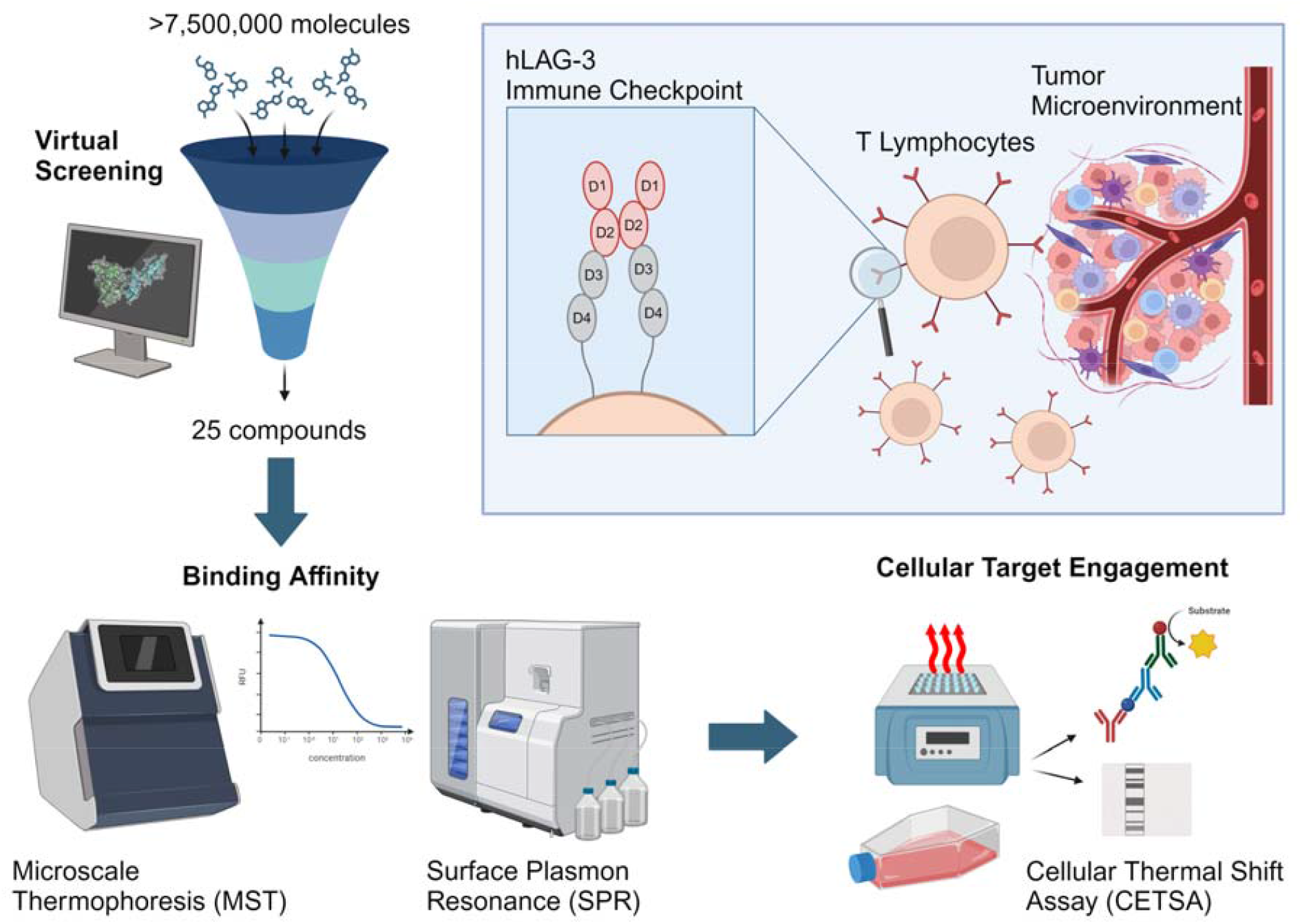
Overall workflow of our study. The immune checkpoint LAG-3 plays a role in modulating the imm response in the tumor microenvironment. Potential LAG-3 ligands were selected in a virtual screening. Then, their binding affinity was evaluated using Microscale Thermophoresis (MST) and Surface Plasmon Resonance (SPR). A Cellular Thermal Shift Assay (CETSA) was used to assess their potential to engage LAG-3 on a cellular level. Figure

In order to identify potential small molecule binding sites of LAG-3 (PDB ID: 7TZH, domains 3-4^16^), we per-formed molecular dynamics (MD) simulations. The t**r**ajectories served as input for PyRod^17^, an in-house developed tool that tracks the movement of water molecules and thus identifies interactions on the protein surface for bi**n**ding small molecules. We identified a lipophilic canyon that could serve as a binding site as previously reported f**o**r Ig-like proteins^14,18^ (Figures 2A–C). Using this informatio**n**, we developed a 3D pharmacophore using LigandScout^19^ (Figures 2D, F). It contains a hydrogen bond acceptor dir**e**cted towards the backbone of the residue F380, two hydrogen bond donors directed towards the backbone of C36**9** and the side chain of E370 with an additional positive ioni**z**able feature also showing an interaction with the side ch**a**in of E370. Additionally, four hydrophobic features were i**d**entified in close proximity to P396, V374, and F380 (Figure 2A). This 3D pharmacophore model was used for a virtual screening campaign of 7.5 million molecules, resulting in 6,696 hits. These hits were then filtered by mole**c**ular weight (MW) to obtain lead-like molecules (MW ≥30**0** Da). Next, unwanted structures and pan-assay interfe**r**ence compounds (PAINS)^20^ were removed resulting in **4**,738 remaining molecules. These were then docked int**o** the lipophilic canyon of LAG-3. The docking poses were **m**inimized using MMFF94^21-25^ to ensure geometrically plausible binding poses. The minimized poses were rescored against the 3D pharmacophore to verify that the desired int**e**ractions were indeed fulfilled. The top-ranked 500 do**c**king poses were selected for visual inspection. During visual inspection, ligand-protein interactions, shape complementarity, and ligand conformation were carefully evaluated, resulting in 17 compounds being selected for experimental testing (Figure S2, Table S1).

**Figure 2.**
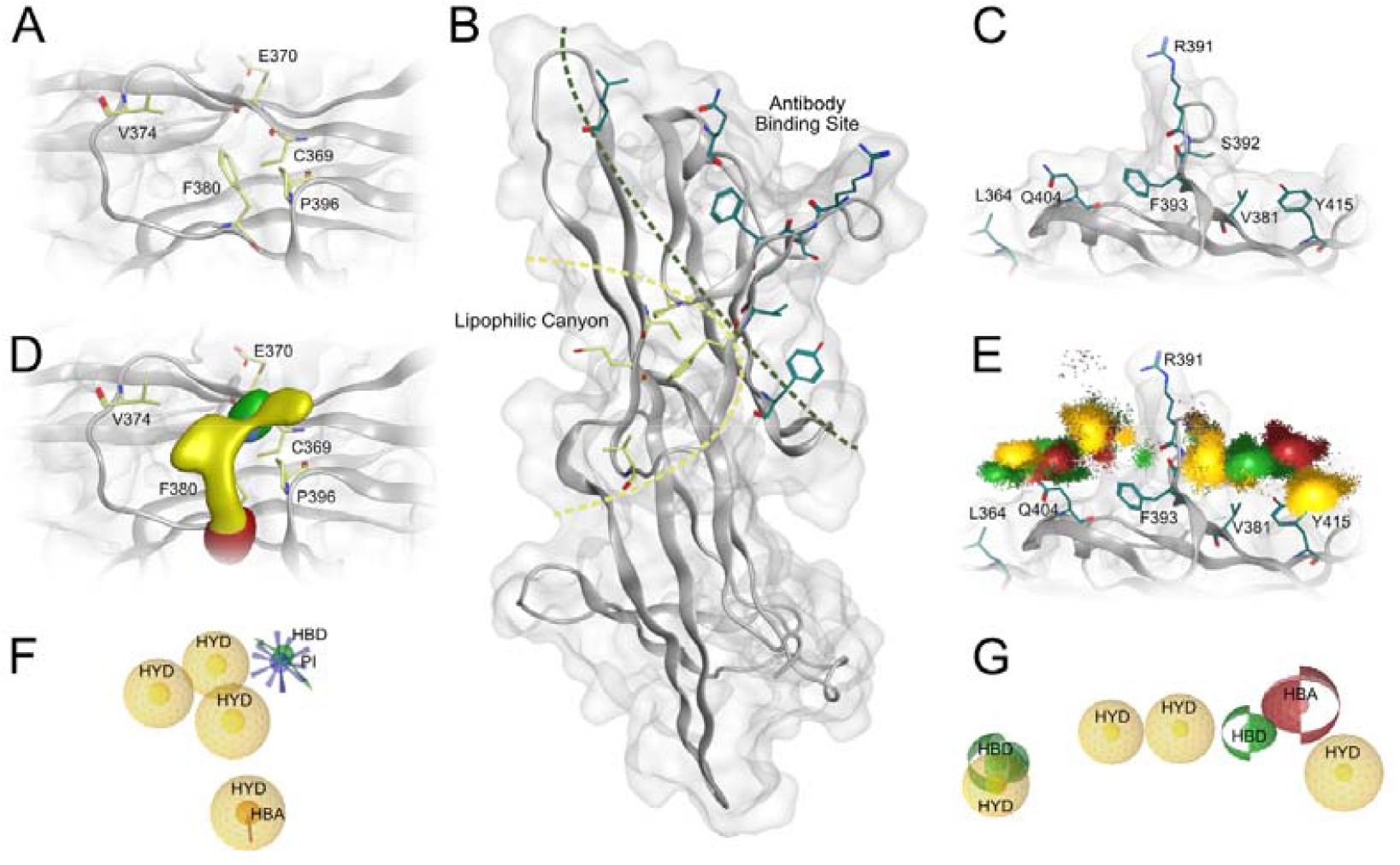
⍰ Identified binding pockets of LAG-3 and 3D pharmacophores for virtual screening. **A**. Lipophilic canyon pocket. **B**. Global view on the LAG-3 ectodomain (color code: grey ribbon and surfaceLAG-3). C. Monoclonal antibody (mAb) binding pocket (color code: yellow stickscarbon atoms of the lipophilic canyon residues; green stickscarbon atoms of the mAb binding pocket residues). **D**. Dynamic molecular interaction fields generated by PyRod. **E**. Dynamic pharmacophore interaction clouds (dynophores). **F**. 3D pharmacophore for binding to the lipophilic canyon. **G**. 3D pharmacophore for binding to the mAb binding pocket. Color code: yellow clouds and spheres - lipophilic contacts; green clouds, arrows, and spheres - hydrogen bond donor; red clouds, arrows, and spheres - hydrogen bond acceptor; purple cloud and star - cationic interaction.

In a second approach, we performed MD simulations of the mAb in complex with LAG-3 (PDB ID: 7TZH^16^) with further analysis of the interactions between the mAb and LAG-3 through the application of Dynophores.^26^ This allowed us to generate a 3D pharmacophore model (Figures 2E, G) consisting of a hydrogen bond acceptor pointing to the amide backbone of S392, a hydrogen bond donor oriented towards the carbonyl oxygen in the backbone of S392, and a second hydrogen bond donor situated adjacent to the side chain of Q404. Four additional hydrophobic features are located next to the sided chains of residues L364, V381, R391, F393, and Y415 (Figure 2C). This 3D pharmacophore was employed in a virtual screening campaign conducted using LigandScout^19^, yielding 97,698 hits, which were filtered according to their MW keeping leadlike molecules. To establish an additional ionic interaction with R391, only negatively charged molecules were retained. We hypothesized that an electrostatic attraction between the small molecule and the binding site might increase the chances of finding a LAG-3 binder^27^. Finally, the unwanted structures and PAINS^20^ were removed, resulting in remaining 973 compounds. These were docked into the binding interface of the mAb with LAG-3 and subsequently minimized using MMFF94^21-25^ to obtain geometrically plausible binding modes. The docking poses were re-scored against the 3D pharmacophore. The 500 topranked docking poses according to the pharmacophore-fit score underwent visual inspection. We selected eight compounds for experimental testing based on the fulfilling pharmacophoric interactions with the protein, ligand conformation, and shape complementarity (Figure S1, Table S1).

We screened a total of 25 compounds that derived from the virtual screening (8 ligand binding site hits, 17 lipophilic canyon hits, see Figure S1-S2) using microscale thermophoresis (MST). Compounds with a signal-to-background ratio ≥5 were considered binders. The MST screening revealed six compounds as binders at 200 µM ligand concentration, two mAb binding site as well as three canyon pocket hits (Figure 3). Next, we performed binding affinity experiments with the same screening platform to determine K_D_ values. Here, we identified two of the binders as dose-dependent ligands of His-tagged LAG-3, compound 3 (mAb / ligand binding site) and compound 25 (canyon pocket) with K_D_ values in the low micromolar range (1.23 ± 0.71 µM and 5.83 ± 3.03 µM, respectively) (Figure 3).

**Figure 3.**
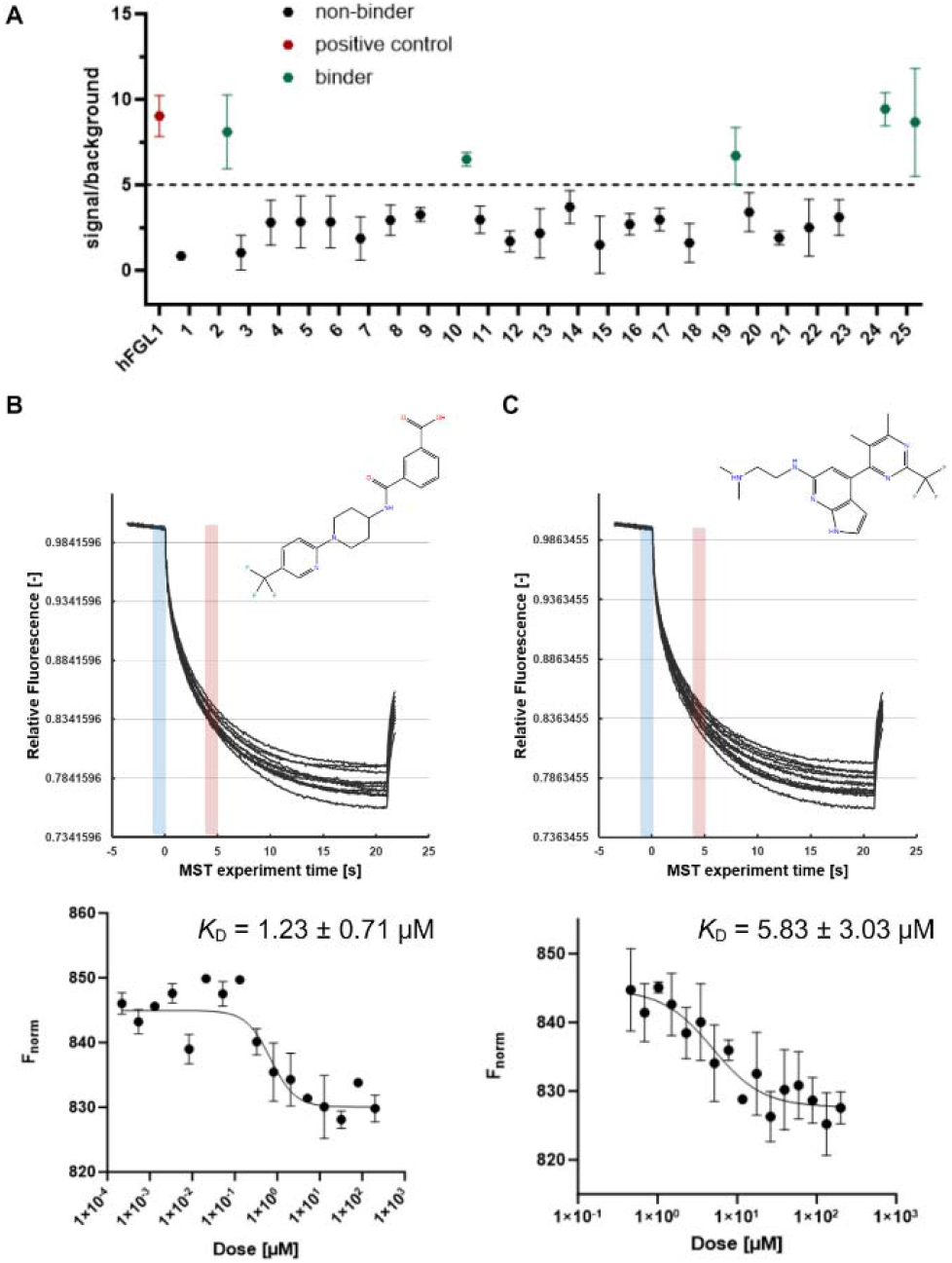
MST screening and binding affinity results for virtual screening hits. **A**. Overview of MST screening results at 200 µM, n = 3. Compounds with a signal-to-background ratio ≥5 were considered binders, FGL1 was used as a positive control. **B**. Structure of compound 3 (mAb / ligand binding site) and MST traces for LAG-3, His tag, incubated with compound **3** including graph for *K*D determination, 2.5-fold dilution series starting at 200 µM, three independent experiments, shown as means ± standard deviations. **C**. Structure of compound **25** (lipophilic canyon) and MST traces for LAG-3, His tag, incubated with compound **25** including graph for *K_D_* determination, 1.5-fold dilution series starting at 200 µM, three independent experiments, displayed as means ± standard deviations. Plots created with GraphPad Prism 10; figure composed with BioRender.

To confirm these latter results, we measured the binding affinity of both complexes (compound 3/LAG-3 and compound 25/LAG-3) by SPR^28^ and confirmed that both of them bound to target protein LAG-3 with *K*_D_ values equal to 8.57 ± 4.57 µM and 2.94 ± 1.12 µM, respectively (Figures 4A, B). Thus, the orthogonal validation of these two first-in-class small molecule LAG-3 binders confirms that the PyRod-based virtual screening approach is successful on discovering small molecule binders for different immune checkpoints and that the Dynophore-based might have the potential to predict immune checkpoint small molecule binders.

**Figure 4.**
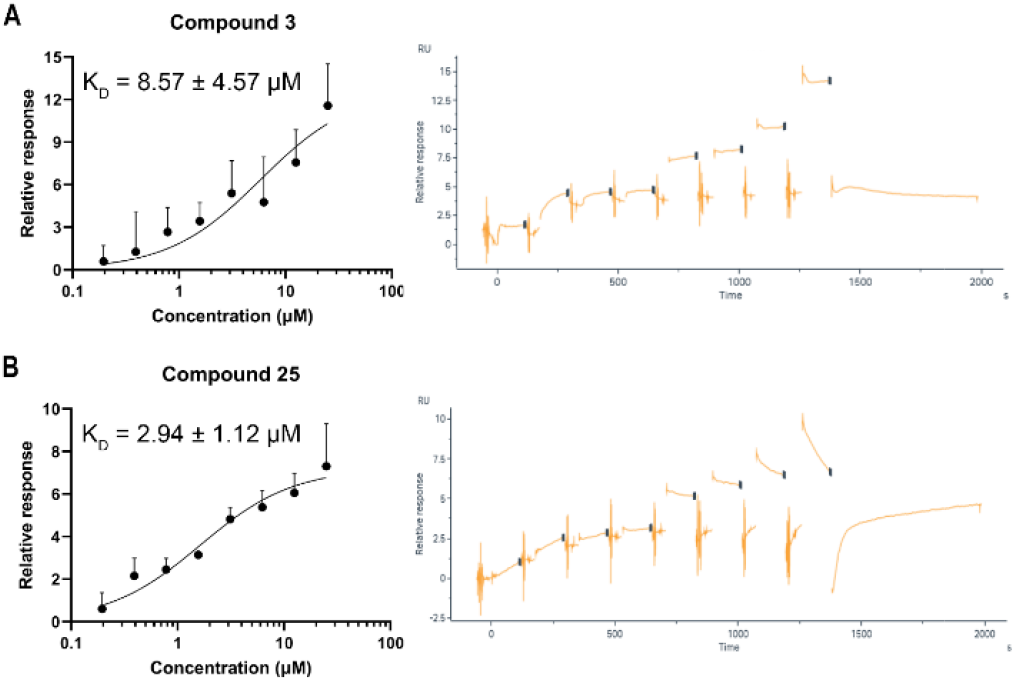
SPR binding affinity results for compounds 3 and 25. **A**. Steady state affinity of compound **3** (25 µM to 195.31 nM) binding to LAG-3 by SPR. **B**. Steady state affinity of compound 25 (25 µM to 195.31 nM) binding to L**A**G-3 by SPR. Graph shows the results of three and four independent experiments, respectively. The data are presented as means ± standard deviations.

Putative binding modes for compounds **3** and **25** to LAG-3 are shown in Figure 5. Compound **25** interacts with several nonpolar amino acids in the lipophilic canyon like V374 and F380. The amine moiety form hydrogen bonds with the carboxyl group of E370 (Figure 5A). Compound 3 can also form hydrogen bonds with S392 and R391 in the mAb binding site. Additionally, there is potential for π-π stacking interactions with F393 (Figure 5B).

**Figure 5.**
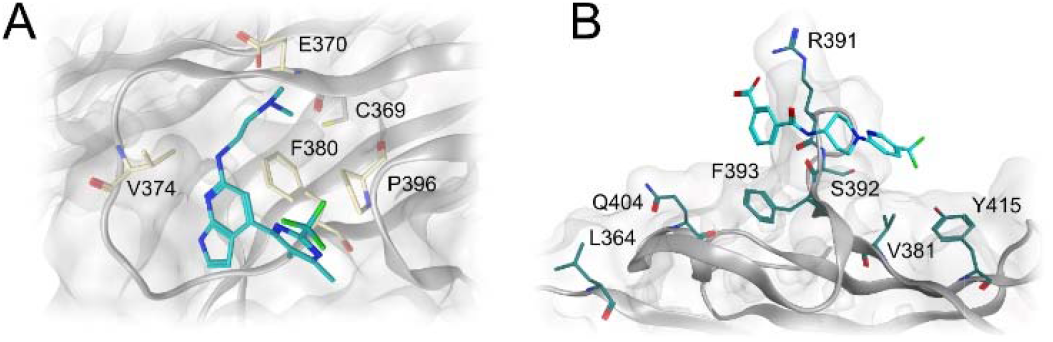
Putative binding modes of the LAG-3 binders. **A**. Binding mode of compound **25** to the lipophilic canyon. **B**. Binding mode of compound **3** to the mAb binding pocket (color code: grey ribbon and surface-LAG-3; yellow sticks-carbon atoms of the lipophilic canyon residues; green sticks-carbon atoms of the mAb binding pocket residues; blue and cyan sticks-carbon atoms of the LAG-binders).

The cellular thermal shift assay (CETSA) was first established by Martinez Molina and co-workers.^29^ Based on lig- and-induced thermal stabilization of the target pr**o**tein, ligand binding to the target in a cellular environment can be evaluated.^29, 30^ This is crucial for drug development since a drug’s *in vivo* efficacy does not only rely on the binding affinity to the target protein, but also on pharmacokinetic properties that limit the amount of ligand that can reach it.^31^ CETSA enables direct monitoring of proteinligand interactions on a cellular level. So far, there have not been any efforts to establish this platform for LAG-3. Here, we show the results of a modified protocol that we tailored to LAG-3, using two validated virtual screening hits as ligands (compounds **3** and **25**). Our model system consisted of Raji-hLAG-3 cells; a human B lymphocyte-derived cell line engineered to stably overexpress hLAG-3. Cells are incubated with the ligands of interest at a single concentration (e.g., 100 µM), harvested, tested for cell viability, and then aliquoted. Each cell suspension aliquot is heated to different temperature endpoints for three minutes. Ming and co-workers report a melting temperature of 47.4 °C for hLAG-3, so we chose temperature endpoints between 46 and 74 °C for the experiment.^13^ After this brief heat treatment, the cells are lysed, and the soluble protein in the lysates can be analyzed. Ligand binding can be observed through stabilization of hLAG-3, which implies that at higher temperatures more protein should remain in solution than in controls without ligand.

First, the tested compounds did not influence cell viability as shown in Figure S4 after a one-hour incubation with Raji-hLAG-3 cells. Since hLAG-3 is a membrane-bound protein and only small amounts of a soluble form are released into serum,^32^ we adapted the cell lysis procedures from the original protocol developed for cytosolic proteins.^30^ To solubilize the membrane-bound hLAG-3 fraction, we added a nonionic detergent to the lysis buffer as described in the literature (Figure S3A).^33, 34^ We then used both ELISA and Western blots to analyze the soluble protein in the cell lysates. Western blots appeared to be more sensitive to small amounts of hLAG-3, whereas ELISA could provide direct quantitative and more robust results. For samples incubated with compound **3**, we observed a moderate stabilization of hLAG-3 at 50 °C with both methods (Figure 6**Error! Reference source not found**.) and a stabilizing effect at 46 °C using ELISA. In contrast, hLAG-3 levels in the DMSO controls decreased rapidly at temperatures higher than 46 °C, matching the reported melting temperature (47.4 °C).^13^ This suggests that compound 3 is able to engage its target hLAG-3 in a cellular environment, making it a promising candidate for future studies. Compound **25**, on the other hand, only displayed minor and inconsistent effects. The ELISA analysis revealed a slight target stabilization at 74 °C with high standard deviations (Figure 6**Error! Reference source not found.A**), but a destabilization at 50 °C and 54 °C. The Western blots, however, showed that it stabilizes hLAG-3 at 50 °C and 54 °C (Figure 6**Error! Reference source not found.B**). Due to the inconsistent results including high standard deviations for compound 25, a lipophilic canyon binder, there is no clear evidence for target engagement in a cellular setting. A possible reason for this could be that the antibody binding surface is more accessible in a cellular environment than the lipophilic canyon, thus impeding the binding of **25** to hLAG-3. Other than that, the compounds’ physicochemical properties, such as lipophilicity, solubility, or stability in culture media, could influence this outcome.

**Figure 6.**
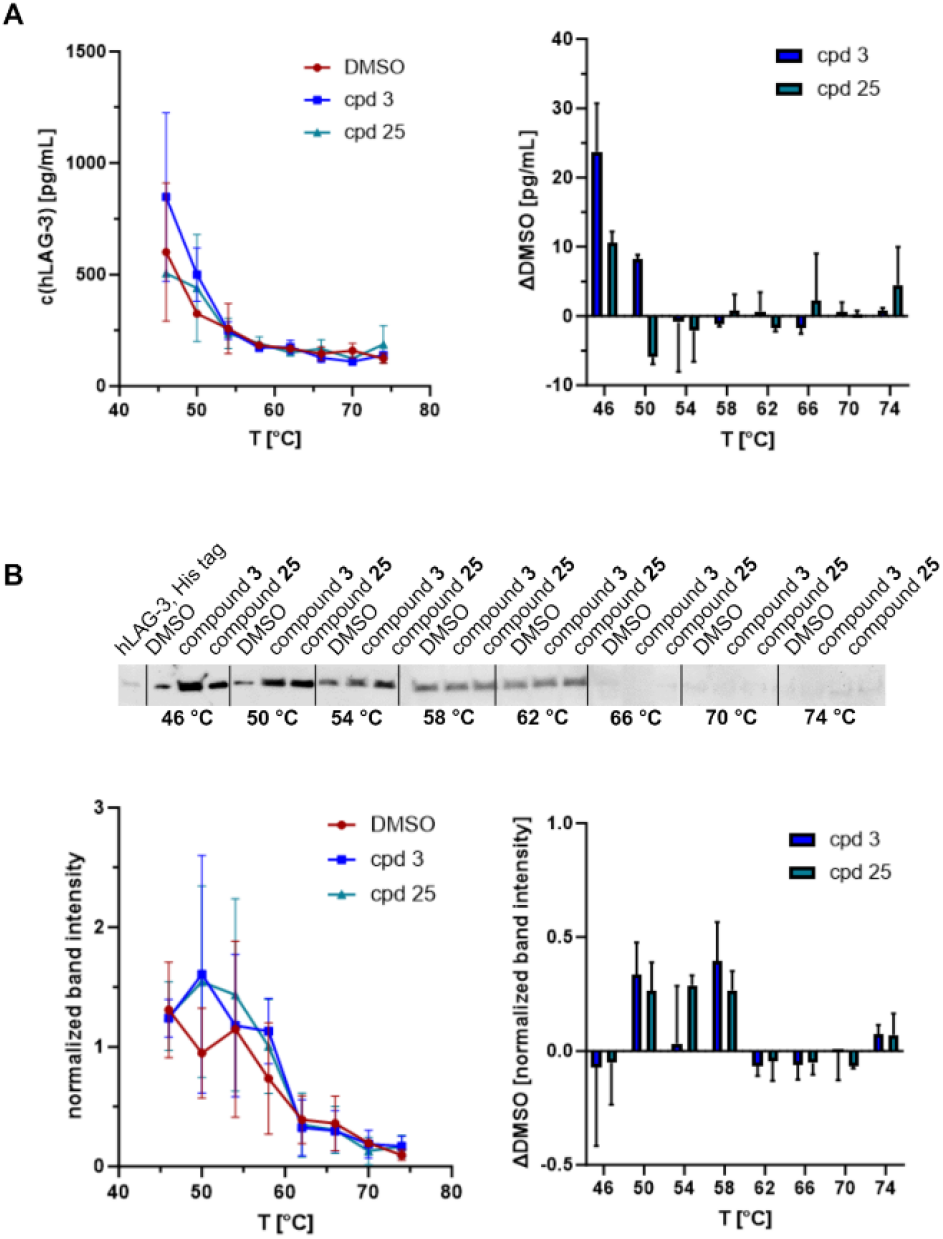
CETSA results for compounds 3 and 25 tested at 100 µM, 0.2% DMSO. **A**. ELISA analysis for all samples and controls, hLAG-3 concentration vs temperature plot (left) and bar plot including the difference of ligand samples vs DMSO controls (right). **B**. Western blot analysis for all samples and controls with one exemplified blot, recombinant hLAG-3, His tag was used as a reference. LAG-3 concentration is presented as normalized band intensity (normalized to total protein), hLAG-3 concentration vs temperature plot (left) and bar plot including the difference of ligand samples vs DMSO controls (right). All graphs show the results of three independent experiments with the data presented as means ± standard deviations. All plots were created with GraphPad Prism 10. Figure composed with BioRender.

Developing small molecule-based checkpoint inhibitors remains a challenging task in drug discovery. Howeve**r**, MD simulation-supported 3D pharmacophore screening has proven to be a powerful tool in identifying binders for novel binding pockets that have not been previously reported in literature. Both the mAb binding site hit (compound **3**, Figure 5B) and the canyon binding site hit (compound **25**, Figure 5A) are single-digit micromolar LAG-3 binders verified by MST and SPR studies. Additionally, we investigated whether the ligands can bind to hLAG-3 on a cellular level using a cellular thermal shift assay with a Raji-hLAG-3 cell model. Compound **3**, an mAb binding site hit, showed a stabilizing effect on hLAG-3 without affecting cell viability. For compound **25**, a potential lipophilic canyon binder, we did not observe protein stabilization. As a new scaffold that binds hLAG-3 in the low micromolar range (*K*_D_ (MST) = 1.2 ± 0.71 µM, *K*_D_ (SPR) = 8.6 ± 4.6 µM) and can also engage its target in a cellular environment, compound **3** emerges as a promising small molecule-based ligand for LAG-3 that should be investigated in future studies. These studies could include cellular and in vivo screening to develop next-generation cancer diagnostics and therapies based on targeting LAG-3.

## Supporting Information

The Supporting Information is available free of charge on the ACS Publications website.

Experimental procedures, virtual screening hits, Western blots, and cell viability studies (PDF)

## AUTHOR INFORMATION

### Author Contributions

The manuscript was written through contributions of all authors. All authors have given approval to the final version of the manuscript.

## ABBREVIATIONS

ADME: absorption, distribution, metabolism, excretion
CETSA: cellular thermal shift assay
CTLA-4: anti-cytotoxic T lymphocyte associated protein 4
ELISA: enzyme-linked immunosorbent assay
FGL1: fibirinogen-like protein 1
hLAG-3: lymphocyte-activation gene 3
ICOS: inducible T cell co-stimulator
mAb: monoclonal antibody
MD: molecular dynamics
MHC: major histocompatibility complex
MST: microscale thermophoresis
MW: molecular weight
PAINS: pan-assay interference compounds
PD-L1: programmed death-ligand 1
SAR: structure-activity relationship
SPR: surface plasmon resonance
TIGIT: anti-T cell immunoreceptor with immunoglobulin and ITIM domain
VISTA: V-domain Ig suppressor of T cell activation

## Notes

The authors declare no competing financial interests.

## ACKNOWLEDGMENTS

We gratefully acknowledge financial support from the ELSA U. Pardee Foundation (Award ID: 2022-215)

## Insert Table of Contents artwork here

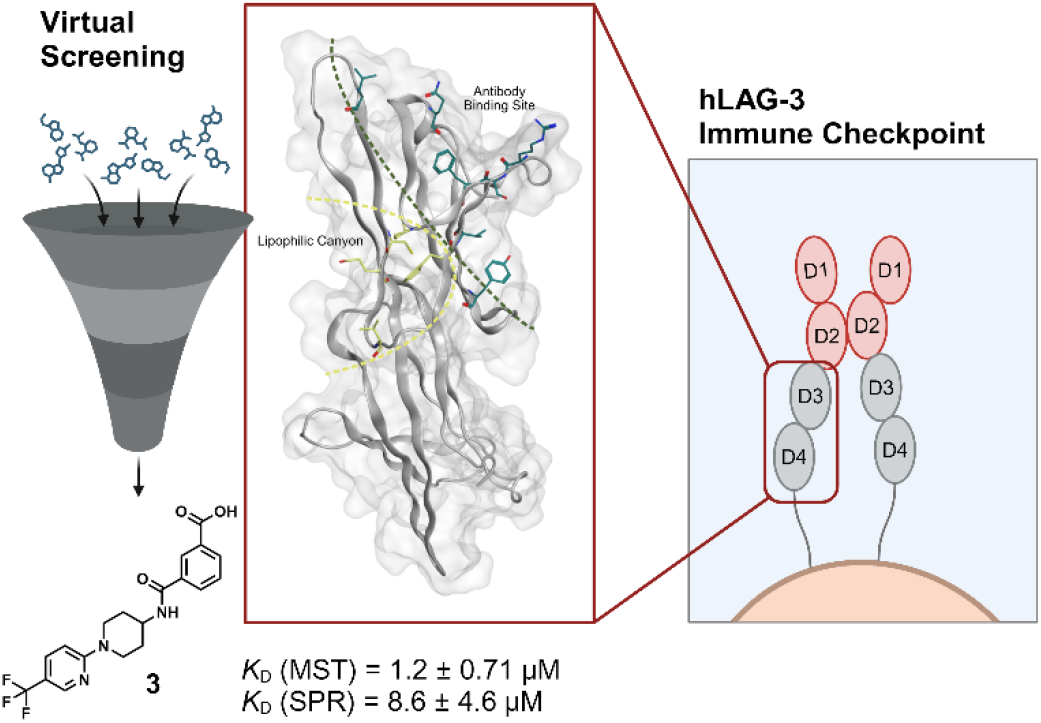

## Notes

### Competing Interest Statement

The authors have declared no competing interest.

## REFERENCES

(1) Tawbi, H. A.; Schadendorf, D.; Lipson, E. J.; Ascierto, P. A.; Matamala, L.; Castillo Gutierrez, E.; Rutkowski, P.; Gogas, H. J.; Lao, C. D.; De Menezes, J. J.; et al. Relatlimab and Nivolumab versus Nivolumab in Untreated Advanced Melanoma. N Engl J Med 2022, 386 (1), 24–34. DOI: 10.1056/NEJMoa2109970.

(2) Hodi, F. S.; O’Day, S. J.; McDermott, D. F.; Weber, R. W.; Sosman, J. A.; Haanen, J. B.; Gonzalez, R.; Robert, C.; Schadendorf, D.; Hassel, J. C.; et al. Improved survival with ipilimumab in patients with metastatic melanoma. N Engl J Med 2010, 363 (8), 711–723. DOI: 10.1056/NEJMoa1003466 From NLM Medline.

(3) Rittmeyer, A.; Barlesi, F.; Waterkamp, D.; Park, K.; Ciardiello, F.; von Pawel, J.; Gadgeel, S. M.; Hida, T.; Kowalski, D. M.; Dols, M. C.; et al. Atezolizumab versus docetaxel in patients with previously treated non-small-cell lung cancer (OAK): a phase 3, open-label, multicentre randomised controlled trial. Lancet 2017, 389 (10066), 255–265. DOI: 10.1016/S0140-6736(16)32517-X From NLM Medline.

(4) Robert, C.; Schachter, J.; Long, G. V.; Arance, A.; Grob, J. J.; Mortier, L.; Daud, A.; Carlino, M. S.; McNeil, C.; Lotem, M.; et al. Pembrolizumab versus Ipilimumab in Advanced Melanoma. N Engl J Med 2015, 372 (26), 2521–2532. DOI: 10.1056/NEJMoa1503093 From NLM Medline.

(5) Robert, C.; Thomas, L.; Bondarenko, I.; O’Day, S.; Weber, J.; Garbe, C.; Lebbe, C.; Baurain, J. F.; Testori, A.; Grob, J. J.; et al. Ipilimumab plus dacarbazine for previously untreated metastatic melanoma. N Engl J Med 2011, 364 (26), 2517–2526. DOI: 10.1056/NEJMoa1104621 From NLM Medline.

(6) Schachter, J.; Ribas, A.; Long, G. V.; Arance, A.; Grob, J. J.; Mortier, L.; Daud, A.; Carlino, M. S.; McNeil, C.; Lotem, M.; et al. Pembrolizumab versus ipilimumab for advanced melanoma: final overall survival results of a multicentre, randomised, open-label phase 3 study (KEYNOTE-006). Lancet 2017, 390 (10105), 1853–1862. DOI: 10.1016/S0140-6736(17)31601-X From NLM Medline.

(7) Leach, D. R.; Krummel, M. F.; Allison, J. P. Enhancement of antitumor immunity by CTLA-4 blockade. Science 1996, 271 (5256), 1734–1736. DOI: 10.1126/science.271.5256.1734 From NLM Medline.

(8) https://clinicaltrials.gov/. (accessed 2024 05/15).

(9) Pardoll, D. M. The blockade of immune checkpoints in cancer immunotherapy. Nat Rev Cancer 2012, 12 (4), 252–264. DOI: 10.1038/nrc3239 From NLM Medline.

(10) Imai, K.; Takaoka, A. Comparing antibody and small-molecule therapies for cancer. Nat Rev Cancer 2006, 6 (9), 714–727. DOI: 10.1038/nrc1913 From NLM Medline.

(11) Abdel-Rahman, S. A.; Rehman, A. U.; Gabr, M. T. Discovery of First-in-Class Small Molecule Inhibitors of Lymphocyte Activation Gene 3 (LAG-3). ACS Med Chem Lett 2023, 14 (5), 629–635. DOI: 10.1021/acsmedchemlett.3c00054.

(12) Gabr, M. T.; Gambhir, S. S. Discovery and Optimization of Small-Molecule Ligands for V-Domain Ig Suppressor of T-Cell Activation (VISTA). J Am Chem Soc 2020, 142 (38), 16194–16198. DOI: 10.1021/jacs.0c07276 From NLM Medline.

(13) Ming, Q.; Celias, D. P.; Wu, C.; Cole, A. R.; Singh, S.; Mason, ; Dong, S.; Tran, T. H.; Amarasinghe, G. K.; Ruffell, B.; et al. LAG3 ectodomain structure reveals functional interfaces for ligand and antibody recognition. Nat Immunol 2022, 23 (7), 1031–1041. DOI: 10.1038/s41590-022-01238-7.

(14) Calvo-Barreiro, L.; Talagayev, V.; Pach, S.; Abdel-Rahman, S. A.; Wolber, G.; Gabr, M. T. Discovery of ICOS-Targeted Small Molecules Using Pharmacophore-Based Screening. ChemMedChem 2023, 18 (23), e202300305. DOI: 10.1002/cmdc.202300305.

(15) Rujas, E.; Cui, H.; Sicard, T.; Semesi, A.; Julien, J. P. Structural characterization of the ICOS/ICOS-L immune complex reveals high molecular mimicry by therapeutic antibodies. Nat Commun 2020, 11 (1), 5066. DOI: 10.1038/s41467-020-18828-4 From NLM Medline.

(16) Ming, Q.; Celias, D. P.; Wu, C.; Cole, A. R.; Singh, S.; Mason, C.; Dong, S.; Tran, T. H.; Amarasinghe, G. K.; Ruffell, B.; et al. LAG3 Ectodomain Structure Reveals Functional Interfaces for Ligand and Antibody Recognition. Nat. Immunol. 2022, 23 (7), 1031–1041. DOI: 10.1038/s41590-022-01238-7.

(17) Schaller, D.; Pach, S.; Wolber, G. PyRod: Tracing Water Molecules in Molecular Dynamics Simulations. J. Chem. Inf. Model. 2019, 59 (6), 2818–2829. DOI: 10.1021/acs.jcim.9b00281.

(18) Abdel-Rahman, S. A.; Talagayev, V.; Pach, S.; Wolber, G.; Gabr, M. T. Discovery of Small-Molecule TIM-3 Inhibitors for Acute Myeloid Leukemia Using Pharmacophore-Based Virtual Screening. J. Med. Chem. 2023, 66 (16), 11464–11475. DOI: 10.1021/acs.jmedchem.3c00960.

(19) Wolber, G.; Langer, T. LigandScout: 3-D Pharmacophores Derived from Protein-Bound Ligands and Their Use as Virtual Screening Filters. J. Chem. Inf. Model. 2005, 45 (1), 160–169. DOI: 10.1021/ci049885e.

(20) Baell, J. B.; Holloway, G. A. New Substructure Filters for Removal of Pan Assay Interference Compounds (PAINS) From Screening Libraries and for Their Exclusion in Bioassays. J. Med. Chem. 2010, 53 (7), 2719–2740. DOI: 10.1021/jm901137j.

(21) Halgren, T. A. Merck Molecular Force Field. I. Basis, Form, Scope, Parameterization, and Performance of MMFF94. J. Comput. Chem. 1996, 17 (5-6), 490–519. DOI: 10.1002/(sici)1096-987x(199604)17:5/6<490::Aid-jcc1>3.0.Co;2-p.

(22) Halgren, T. A. Merck Molecular Force Field. II. MMFF94 Van Der Waals and Electrostatic Parameters for Intermolecular Interactions. J. Comput. Chem. 1996, 17 (5-6), 520–552. DOI: 10.1002/(sici)1096-987x(199604)17:5/6<520::Aid-jcc2>3.0.Co;2-w.

(23) Halgren, T. A. Merck Molecular Force Field. III. Molecular Geometries and Vibrational Frequencies for MMFF94. J. Comput. Chem. 1996, 17 (5-6), 553–586. DOI: 10.1002/(sici)1096-987x(199604)17:5/6<553::Aid-jcc3>3.0.Co;2-t.

(24) Halgren, T. A.; Nachbar, R. B. Merck Molecular Force Field. IV. Conformational Energies and Geometries for MMFF94. J. Comput. Chem. 1996, 17 (5-6), 587–615. DOI: 10.1002/(sici)1096-987x(199604)17:5/6<587::Aid-jcc4>3.0.Co;2-q.

(25) Halgren, T. A. Merck Molecular Force Field. V. Extension of MMFF94 Using Experimental Data, Additional Computational Data, and Empirical Rules. J. Comput. Chem. 1996, 17 (5-6), 616–641. DOI: 10.1002/(sici)1096-987x(199604)17:5/6<616::Aid-jcc5>3.0.Co;2-x.

(26) Bock, A.; Bermudez, M.; Krebs, F.; Matera, C.; Chirinda, B.; Sydow, D.; Dallanoce, C.; Holzgrabe, U.; De Amici, M.; Lohse, M. J.; et al. Ligand Binding Ensembles Determine Graded Agonist Efficacies at a G Protein-Coupled Receptor. J. Biol. Chem. 2016, 291 (31), 16375–16389. DOI: 10.1074/jbc.M116.735431.

(27) Honig, B.; Nicholls, A. Classical electrostatics in biology and chemistry. Science 1995, 268 (5214), 1144–1149. DOI: 10.1126/science.7761829.

(28) Fang, Y. Ligand-receptor interaction platforms and their applications for drug discovery. Expert Opin Drug Discov 2012, 7 (10), 969–988. DOI: 10.1517/17460441.2012.715631.

(29) Martinez Molina, D.; Jafari, R.; Ignatushchenko, M.; Seki, T.; Larsson, E. A.; Dan, C.; Sreekumar, L.; Cao, Y.; Nordlund, P. Monitoring drug target engagement in cells and tissues using the cellular thermal shift assay. Science 2013, 341 (6141), 84–87. DOI: 10.1126/science.1233606.

(30) Jafari, R.; Almqvist, H.; Axelsson, H.; Ignatushchenko, M.; Lundback, T.; Nordlund, P.; Martinez Molina, D. The cellular thermal shift assay for evaluating drug target interactions in cells. Nat Protoc 2014, 9 (9), 2100–2122. DOI: 10.1038/nprot.2014.138.

(31) Ruiz-Garcia, A.; Bermejo, M.; Moss, A.; Casabo, V. G. Pharmacokinetics in drug discovery. J Pharm Sci 2008, 97 (2), 654–690. DOI: 10.1002/jps.21009.

(32) Triebel, F. LAG-3: a regulator of T-cell and DC responses and its use in therapeutic vaccination. Trends Immunol 2003, 24 (12), 619–622. DOI: 10.1016/j.it.2003.10.001 From NLM Medline.

(33) Kawatkar, A.; Schefter, M.; Hermansson, N. O.; Snijder, A.; Dekker, N.; Brown, D. G.; Lundback, T.; Zhang, A. X.; Castaldi, M. P. CETSA beyond Soluble Targets: a Broad Application to Multipass Transmembrane Proteins. ACS Chem Biol 2019, 14 (9), 1913–1920. DOI: 10.1021/acschembio.9b00399.

(34) Kalxdorf, M.; Gunthner, I.; Becher, I.; Kurzawa, N.; Knecht, S.; Savitski, M. M.; Eberl, H. C.; Bantscheff, M. Cell surface thermal proteome profiling tracks perturbations and drug targets on the plasma membrane. Nat Methods 2021, 18 (1), 84–91. DOI: 10.1038/s41592-020-01022-1 Medline. From NLM

