## Supporting Information for "From Virtual Screens to Cellular Target Engagement: New Small Molecule Ligands for the Immune Checkpoint LAG-3"

*Electronic Supplementary Information*

### Experimental Procedures

#### Virtual Screening

Protein Preparation and Molecular Dynamics Simulations

The structure of human LAG-3 with the highest resolution of 2.43 Å (PDB ID: 7TZH^1^) was prepared: 1) in its complex with a monoclonal antibody (mAb) and 2) apo form after the removal of the antibody. The crystallographic water and 2-acetamido-2-deoxy-beta-D-glucopyranose (NAG) covalently bound to hLAG-3 were kept. Both, mAb-complex and apo hLAG-3, were protonated at pH 7 and temperature of 300 K using Protonate 3D function^2^ implemented in MOE 2019.0101 (Chemical Computing Group, Montreal, Canada).

The mAb complex and apo LAG-3 were prepared for molecular dynamics (MD) simulations using Maestro 11.7 (Schrödinger, LLC, New York, USA). The hydrogen bond network in both systems was optimized at pH 7.0. The protein was placed in an orthorhombic box keeping the edges in a 10 Å distance to the protein surface. The box was filled with TIP4P water model,^3^ sodium, and chloride ions to neutralize the system and achieve isotonic conditions (0.15 M NaCl). The MD simulations were performed in 10 replicates, 10 ns each, generating 2000 frames per replicon. The mAb complex and apo LAG-3 systems were simulated as NPT ensembles keeping the number of particles, pressure, and temperature constant (p= 1.01325 bar, T= 298 K). The temperature was controlled using the Nose-Hoover thermostat^4, 5^ and the pressure using the Martyna-Tobias-Klein method.^6^ All MD simulations were carried out with Desmond in version 2020-4^7^ on RTX 2080Ti graphics processing units (NVIDIA Corporation, Santa Clara, USA).

3D Pharmacophore-based Virtual Screening, Docking, and Hit Selection

To increase the chance of hLAG-3 binding identification, we selected two potential binding pockets: the mAb-binding site and a lipophilic canyon in proximity to the mAb-binding site as reported previously for other Ig-like domains.^8, 9^

The 3D pharmacophore for virtual screening (VS) campaign for the canyon pocket of the apo hLAG-3 form was identified using PyRod 0.7.1.^10^ Briefly, PyRod samples the interaction frequency and geometry between water molecules and protein over the course of MD simulation to propose the most favorable interaction hot spots for a small molecular binder. The interaction sampling was carried out in a 30 Å box centered on the CE1 atom of F380 with a 1 Å grid spacing. To ensure system relaxation, only the last 5 ns of the simulation time were analyzed.^10^ The automatically generated dynamic molecular interaction fields were extracted to obtain a maximum of 20 interaction points per interaction type. The final apo-pharmacophore was constructed using interactions with a minimum score of 30 indicating interaction hot spots.^9^

The 3D pharmacophore for the mAb-binding pocket was identified using Dynophore 0.1.^11^ Briefly, the interactions between hLAG-3 and two mAb loops consisting of residues T60-Y61-Y62 and Y232-Y233-P234 were sampled over the course of the MD simulations. The spatiotemporal interaction clouds (dynamic pharmacophores, dynophores for short) were extracted in an automated fashion. To generate a static 3D pharmacophore model, the interactions with frequencies over 50% of the simulation time were kept. The size of the static pharmacophore features was adjusted manually to the size of the dynophore clouds.

The PyRod (apo) and Dynophore (mAb complex) 3D pharmacophores were used for VS on the MolPort library in version 2020 using LigandScout 4.2.^12, 13^ The MolPort library was prepared using the idbgen program implemented in LigandScout 4.2. Each ligand yielded maximally 25 conformations with an energy window of 15 kcal/mol and root mean square deviation window between the generated conformations of 0.5 Å.

The obtained initial hits were filtered using RDKit^14^ with a set of in-house SMARTS patterns (Daylight Chemical Information Systems Inc., Laguna Niguel, USA) detecting reactive, chemically labile, or non-drug-like moieties as reported previously.^8, 9^ Only lead-like molecules with a molecular weight of over 300 Da were kept. Furthermore, the mAb-LAG-3-complex-based hits were filtered to contain at least one negative charge to establish an additional interaction with the central residue R131. Finally, the filtered set of hits for PyRod (apo) and Dynophore (mAb complex) 3D pharmacophores were docked to their respective binding pockets using Gold 2022^15^ to obtain plausible binding modes. The apo-LAG-3 binding pocket was defined in a sphere of 10 Å radius with a center on the CE1 atom of F380 The mAb-binding pocket was defined in a sphere of 10 Å radius with a center on the OG atom of S392. Each docking was run with 10 genetic algorithm runs per molecule using the ChemPLP scoring function.^16^ The final set of molecules for biophysical evaluation was selected after visual inspection.^17, 18^

#### Compounds

All tested compounds were obtained from commercial vendors in lyophilized format (**Table S1**, **Figure S1**, **Figure S2**). Small molecules were resuspended in 100% DMSO (Cat #D8418, Sigma Aldrich, St. Louis, MO, USA) at 10 mM or 50 mM and stored at –30 °C until use.

#### Microscale Thermophoresis (MST)

The Protein Labeling Kit RED-NHS 2^nd^ Generation (Cat #MO-L011, NanoTemper Technologies, Munich, Germany) was used for the labeling of the human LAG-3 protein, His tag (Cat # 16498-H08H, Sino Biological, Beijing, China) following manufacturers’ instructions. No buffer exchange was needed before protein labeling and purification. Briefly, a 4.4 µM hLAG-3 protein solution was mixed with a 5.5-fold concentration of RED-NHS 2^nd^ Generation dye in assay buffer and incubated for 30 minutes at room temperature in the dark. The assay buffer was 1x PBS, 0.05% Tween 20, pH 7.4. After protein-dye incubation, LAG-3 purification by removing the remaining free dye was performed following manufacturers’ instructions. The final concentration and the degree of labeling (DOL) of hLAG-3, the number of dye molecules attached to individual protein molecules, were calculated based on absorbance measurements (A_280_ and A_650_, respectively) using a Biotek Synergy Neo2 Reader (Agilent, Santa Clara, CA, USA). Successful labeling was considered if the DOL was comprised between 0.5 and 1. Additionally, the protein fold was confirmed by comparing the melting curve of hLAG-3 before and after protein labeling using the Tycho NT.6 system (NanoTemper Technologies). Finally, labeled hLAG-3 protein was flash-frozen in liquid nitrogen and stored at -80°C until use.

Compounds were diluted in MST buffer (10 mM HEPES, 150 mM NaCl, 0.005% Tween 20, pH 7.4) to the corresponding final concentration in the presence of 2% DMSO. Human FGL1, Fc tag (Cat #FG1-H5258, Acro Biosystems) in MST buffer supplemented with 2% DMSO was used as positive controls while 2% DMSO in MST buffer was used as a negative control.

Freshly thawed labeled LAG-3 stock protein was centrifuged for 10 min at 15,000 rpm and 4°C before each use. Then, the protein was diluted to 100 nM in MST buffer and incubated 1:1 with the corresponding 2-fold concentrated compound or control to a final volume of 20 µL for 10 minutes at room temperature in the dark. All measurements were performed on a NanoTemper Monolith NT.115 (NanoTemper Technologies) using Monolith Standard Capillaries (Cat #MO-K022, NanoTemper Technologies). The system was set to 25 °C, 40% excitation, medium MST power, and samples were measured for 1 second without heating (before infrared laser activation) and during 21 seconds with the infrared laser turned on. Results are displayed as fluorescence values: i) relative fluorescence, fluorescence obtained during the length of the experiment (TRIC) and normalized to the fluorescence obtained at the set temperature; ii) normalized fluorescence (F_norm_, %): ratio between fluorescence values after and before laser activation, as previously described.^19^

In the binding check experiments, four technical replicates were performed per screened small molecule, positive control, or negative control. Small molecules were screened at 200 µM. In the binding affinity experiments, a 16-point serial dilution was prepared for every selected compound starting at 200 µM. Three independent experiments were performed in the single-dose and binding affinity experiments to determine potential binding molecules and affinity constants, respectively. Raw data was plotted in GraphPad Prism 10.

#### Surface Plasmon Resonance (SPR)

The binding analysis was performed on a Biacore^TM^ 8K instrument (Cytiva, Marlborough, MA, USA) at 25°C using 1x PBS-P+ buffer (Cat #28995084, Cytiva) supplemented with 2% DMSO as the running buffer. Biotinylated Human LAG-3 Protein, His, Avitag™ (Cat #LA3-H82E9, Acro Biosystems) was captured on a Series S Sensor Chip CAP at a flow rate of 10 μL/min for 60 seconds yielding an immobilization level of approximately 4,000 RU using the Biotin CAPture Kit (Cytiva). The solutions provided are a Biotin CAPture Reagent, which is a modified streptavidin, for capturing the ligand on to the sensor chip CAP and two regeneration stock solutions for the regeneration of the surface after each run. For binding analysis, serial dilutions of the analyte (25 µM to 195.31 nM, 2-fold dilution, 8 steps) were prepared in running buffer and injected over the chip at a flow rate of 30 μL/min for 120 seconds (association phase). The dissociation phase was set at 10 minutes. Data was obtained using the Biacore 8K Control Software (Cytiva) and analyzed by non-linear curve fitting using a steady-state affinity analysis applying the software Biacore^TM^ Insight Evaluation Software (Cytiva).

#### Cellular Thermal Shift Assay (CETSA)

CETSA procedures were adapted from *Jafari et al* and *Kawatkar et al*.^20, 21^

Cell Culture

Raji-hLAG-3 cells (InvivoGen, San Diego, CA, USA, Cat #raji-hlag3) were grown in 37 °C, 5% CO_2_ using supplemented IMDM (25 mM HEPES, 2 mM l-glutamine, 10% heat-inactivated fetal bovine serum, 100 U/mL-100 µg/mL Pen-Strep, 100 µg/mL Normocin, 10 µg/mL Blasticidin) until they reached a cell density of 2,000,000 cells/mL. The cells were counted, and cell viability was assessed with TrypanBlue. Then, the cells (2,000,000 cells/mL, 15 mL) were incubated with the respective ligands or vehicle (100 µM, 0.2% DMSO) for 1 hour at 37 °C, 5% CO_2_ followed by centrifugation (300 rcf, 3 min, rt). Cell pellets were gently resuspended in 1 mL PBS containing protease inhibitors (Sigma-Aldrich, St. Louis, MO, USA, Cat #P8340, 200-fold dilution) succeeded by cell counting and cell viability assessment with TrypanBlue.Each cell suspension (DMSO control, ligand samples) was divided into ten different tubes, each containing 100 µL cell suspension (approximately 3,000,000 cells per tube).

Temperature Endpoints

Each aliquot was incubated at a different temperature endpoint for 3 min. The endpoints ranged from 46 °C to 74 °C. After heat treatment, the tubes were left at rt for 3 min and then stored at –80 °C until cell lysis.

Cell Lysis

Following incubation at different temperature endpoints, the cells were lysed using PBS pH 7.4, 0.4% NP-40, supplemented with protease inhibitors (Sigma-Aldrich, #P8340, 200-fold dilution) to a final sample volume of 125 µL. After three freeze-thaw cycles (liquid nitrogen, water bath at 37 °C, 3 min each), the cell lysates were centrifuged for 20 min at 11,800 g, 4 °C.

Western Blots

For Western blots, precast SDS-PAGE gels (4–20% Mini-PROTEAN® TGX™ Precast Protein Gels, 15-well, Bio-Rad, Cat #4561096) were loaded with 0.25 ng of recombinant hLAG-3, ECD, His tag (SinoBiological, Cat #16498-H08H), 15 µL of the cell lysate samples in NuPAGE LDS sample buffer (Invitrogen, Cat #NP0007), and 5 µL PageRuler Prestained Protein Ladder (ThermoFisher Scientific, Waltham, MA, USA, Cat #26616). The gels were run at 120 V for
40–50 min in NuPAGE MES SDS running buffer (Invitrogen, Waltham, MA, USA, Cat #NP0002). Proteins were transferred to polyvinylidene difluoride membranes (ThermoFisher Scientific, Cat #LC2002) in transfer buffer (25 mM Tris, 192 mM glycine, 20% w/v methanol, pH 8.3) for 1 hour at 0.35 A, 4 °C. After transfer, the membranes were washed with ultrapure water for 3 x 3 min at rt. Total protein staining was performed using the No-Stain^TM^ Protein Labeling Reagent (ThermoFisher Scientific/Invitrogen, Cat # A44449), following the user guide manual. After total protein detection with a ChemiDoc system (Bio-Rad Laboratories), the membrane was washed with TBST (10mM Tris-HCl, pH 8.0, 150 mM NaCl, 0.1% Tween 20) for 5 min at rt, and non-specific binding was blocked using EveryBlot Blocking Buffer (Bio-Rad, Cat #12010020) for 15 min at rt. The membranes were then incubated with a polyclonal anti-LAG-3 antibody, rabbit / IgG (ThermoFisher Scientific/Proteintech, Cat #16616-1-AP, 1:1,000) with gentle agitation at 4 °C overnight. After washing three times with TBST (5 min each time), the membranes were incubated with a secondary goat-anti rabbit IgG (H+L), HRP reactive antibody (Invitrogen, Cat #65-6120, 1:4,000) for 1 hour at room temperature. Again, the membranes were washed using TBST (3 x 5 min). Protein bands were detected using SuperSignal West Pico PLUS Chemiluminescent Substrate (ThermoFisher Scientific, Cat #34577) with a ChemiDoc system (Bio-Rad Laboratories). Data evaluation was performed using ImageJ and GraphPad Prism 10. All bands were normalized to total protein.

ELISA

The LAG-3 Human ELISA Kit (ThermoFisher Scientific/Invitrogen, Cat #BMS2211) was used according to the manufacturer’s protocol. All samples were analyzed in duplicates. Data evaluation was performed using GraphPad Prism 10.

### Supporting Data

#### Virtual Screening Hits

The following compounds were selected as hits from the virtual screening (**Table S1**).

**Table S1**. Compounds selected for experimental testing.

| **ID** | **Compound Code** | **SMILES** |
| --- | --- | --- |
|  | **mAB/Ligand Binding Site Compounds** | |
| **1** | ZINC1614990570 | Fc1cnc(cc1)CNC(=O)N2C[C@@H](C(F)(F)F)[C@H](C(=O)[O-])C2 |
| **2** | ZINC342017419 | O(c1c(cc(cc1)CNC(=O)Cc2cnc(cc2)C)C(=O)[O-])C |
| **3** | ZINC238207328 | FC(F)(F)c3cnc(N2CCC(NC(=O)c1cc(ccc1)C(=O)[O-])CC2)cc3 |
| **4** | ZINC1597062599 | o1c(c(cc1CNC(=O)c2cnc(OCC)cc2)C)C(=O)[O-] |
| **5** | ZINC124379381 | s1c(nc(c1)C(=O)[O-])CNC(=O)c2cc(OCC)c(cc2)C |
| **6** | ZINC315561042 | s1c(nc(c1C(=O)[O-])C)CNC(=O)c2cc(OC)c(cc2)C |
| **7** | ZINC864123557 | o1c(ccc1CNC(=O)c3cc2OC(Cc2cc3)(C)C)C(=O)[O-] |
| **8** | ZINC1614994075 | Cn1nc(NC(=O)N2C[C@@H](C(=O)O)[C@@H](C(F)(F)F)C2)cc1C1CC1 |
| **9** | ZINC1614989912 | FC(F)(F)[C@@H]2CN(C(=O)NCc1cc(ncc1)C)C[C@H]2C(=O)[O-] |
| **10** | ZINC804232 | S(c1ncccc1C(=O)[O-])CC(=O)Nc3ccc(Oc2ccccc2)cc3 |
|  | **Lipophilic Canyon Compounds** | |
| **11** | ZINC72480158 | s1cnc5c1ccc(n2c(ncc2)c3nn4c(c3)C[NH2+]CC4)c5 |
| **12** | ZINC72151532 | n1(c(ncc1)c2ccc(cc2)[C@H]([NH+](C)C)C)Cc3c(cncc3)C |
| **13** | ZINC72424139 | Fc3c(O)cc(F)c(NC(=O)CCc1nn2c(c1)C[NH2+]CC2)c3 |
| **14** | ZINC12995456 | FC(F)Oc1ccc(cc1)CCNC(=O)c2ccc(cc2)C[NH+](C)C |
| **15** | ZINC72418855 | S(C(F)(F)F)CCn1c(ncc1)c2nn3c(c2)C[NH2+]CCC3 |
| **16** | ZINC71764449 | FC(F)(F)CCc1nc(ncc1)c3cc2c(O[C@H](C[NH3+])C2)cc3 |
| **17** | ZINC91493937 | Fc3cc(c1n[nH]cc1C(=O)NC[C@H]([NH+](C)C)c2oc(cc2)C)ccc3 |
| **18** | ZINC77540416 | Clc1cc(F)c(cc1)C(=O)N3[C@@H](c2ccc(cc2)C[NH+](C)C)CCCC3 |
| **19** | ZINC72454823 | o4nc(OCCn1c(ncc1)c2nn3c(c2)C[NH2+]CCC3)c(n4)C |
| **20** | ZINC72428715 | n34nc(c1n(ccn1)CCc2ccncc2)cc3C[NH2+]CCC4 |
| **21** | ZINC77456094 | O=C(N2[C@@H](c1ccc(cc1)C[NH+](C)C)CCCC2)c3nnccc3 |
| **22** | ZINC263639933 | Clc1c(ncc(c1)C(F)(F)F)c2nc(sc2)NCC[NH+](C)C |
| **23** | ZINC154028682 | S(=O)(=O)(Nc1c(ccc(c1)C[NH+](C)C)C)Cc2cc(F)ccc2 |
| **24** | ZINC65605683 | s1c(nc(c1C(=O)NC[C@@H]([NH+](C)C)C2CC2)C)c3ncccc3 |
| **25** | ZINC72169669 | FC(F)(F)c3nc(c2c1c([nH]cc1)nc(NCC[NH+](C)C)c2)c(c(n3)C)C |
| **26** | ZINC71589666 | Fc3c(c1oc(c(n1)C[NH+](Cc2[nH]ccn2)C)C)ccc(OC)c3 |
| **27** | ZINC124368084 | FC(F)(F)Oc1c(cccc1)CCC(=O)N2[C@H](C[NH2+]CC2)C |
| **28** | ZINC72424937 | s1cnc4c1cc(NC(=O)CCc2nn3c(c2)C[NH2+]CCC3)cc4 |
| **29** | ZINC426501754 | FC(F)(F)c1c(nccc1)c2ccc(cc2)C(=O)N(CC[NH+](C)C)CC |
| **30** | ZINC89938704 | O(c1ncccc1C(=O)N2[C@H](C[NH+](CC2)C)C)c3cnccc3 |

**Figure S1** and **Figure S2** show the compounds that were purchased for experimental testing. The binding modes of compounds **3** and **13** are exemplified in **Figure S3**.

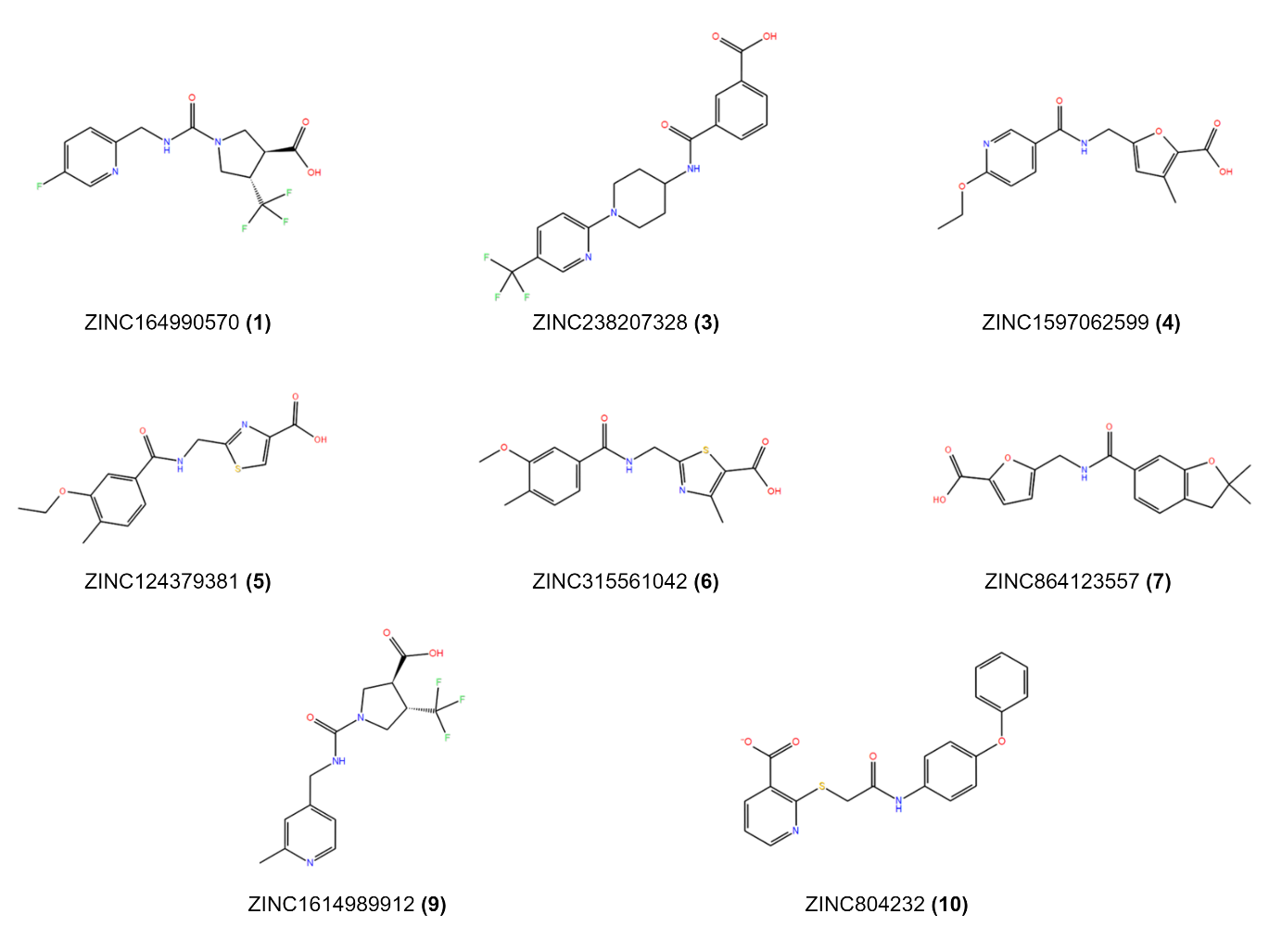

**Figure S1.** Virtual screening hits: antibody interface / ligand binding site. Compounds were purchased from Enamine and ChemBridge.

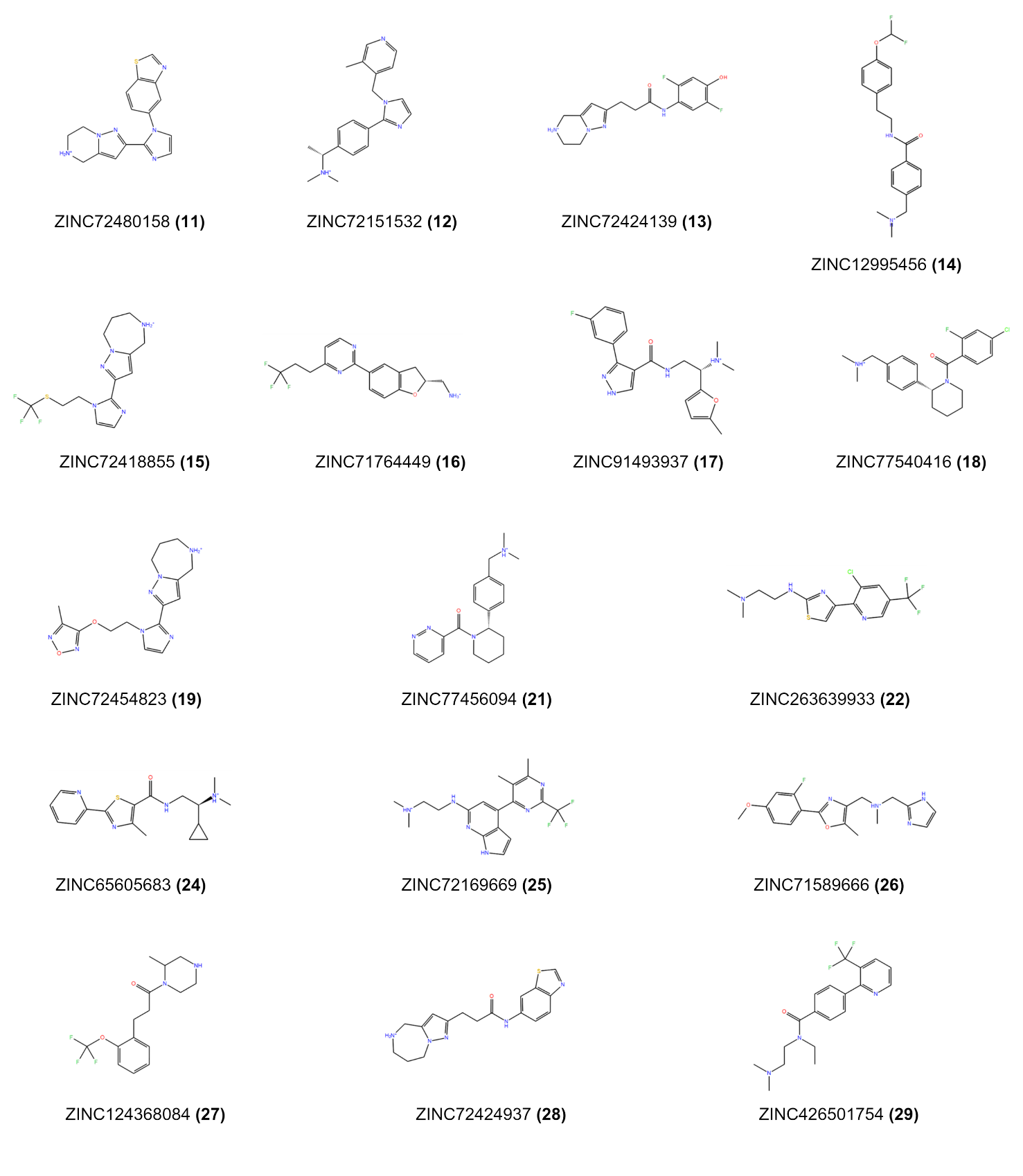

**Figure S2.** Virtual screening hits: lipophilic canyon. Compounds were purchased from Enamine, ChemBridge, and Bionet.

#### Western Blots

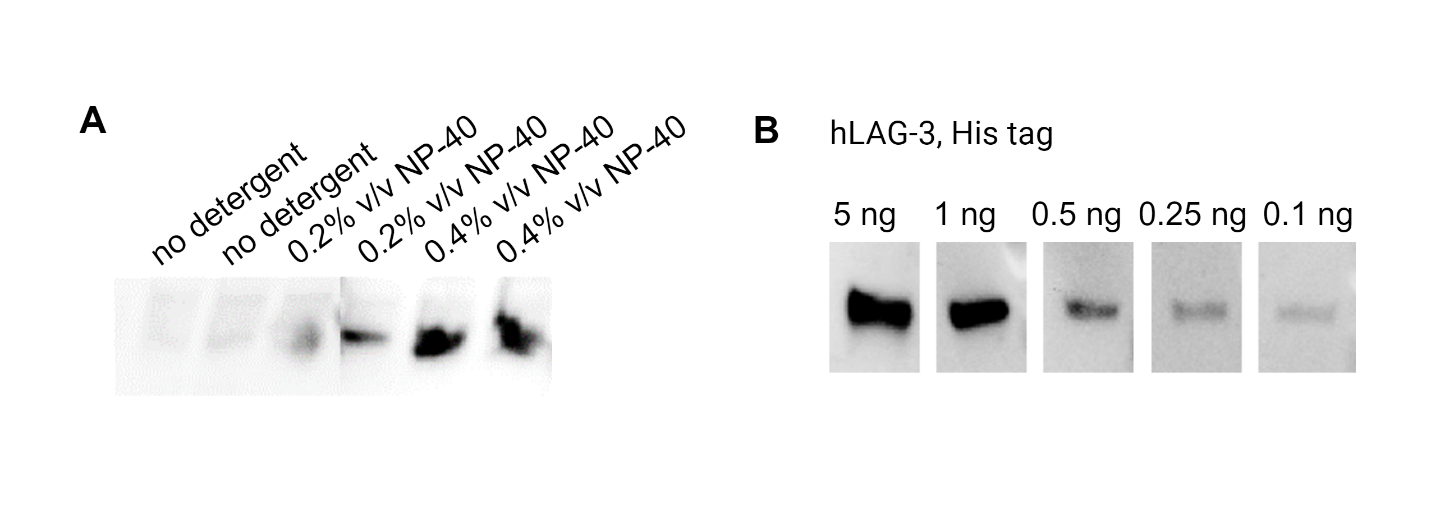

**Figure S3.** Optimization steps. **A.** Optimized cell lysis with addition of a nonionic detergent (NP-40) at different concentrations. Membrane-bound hLAG-3 remains in solution upon adding detergent. **B.** Optimizing the amount of reference protein for the Western blots (hLAG-3, His tag). Bands still visible for 0.1 ng, whereas 5 and 1 ng are already overloaded.

#### Cell Viability

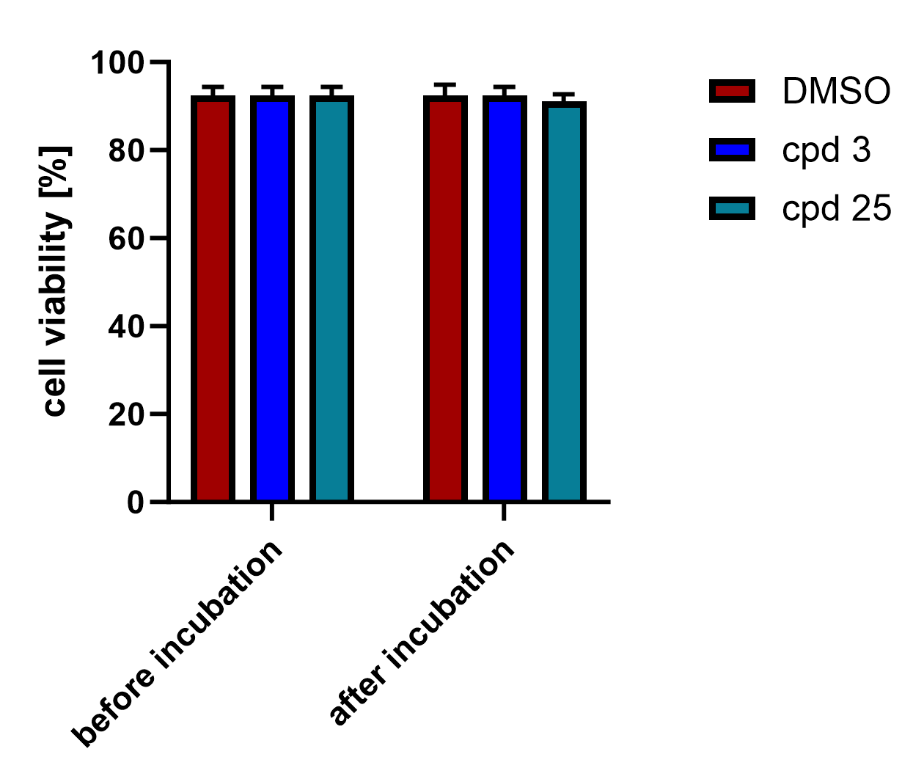

**Figure S4.** Cell viability with TrypanBlue staining before and after 1 h of incubation with DMSO / ligand solutions. No significant effects observed.

### References

(1) Ming, Q.; Celias, D. P.; Wu, C.; Cole, A. R.; Singh, S.; Mason, C.; Dong, S.; Tran, T. H.; Amarasinghe, G. K.; Ruffell, B.; et al. LAG3 ectodomain structure reveals functional interfaces for ligand and antibody recognition. *Nat Immunol* **2022**, *23* (7), 1031-1041. DOI: 10.1038/s41590-022-01238-7.

(2) Labute, P. Protonate3D: assignment of ionization states and hydrogen coordinates to macromolecular structures. *Proteins* **2009**, *75* (1), 187-205. DOI: 10.1002/prot.22234.

(3) Abascal, J. L.; Vega, C. A General Purpose Model for the Condensed Phases of Water: TIP4P/2005. *J. Chem. Phys.* **2005**, *123* (23), 234505. DOI: 10.1063/1.2121687.

(4) Nosé, S. A Molecular Dynamics Method for Simulations in the Canonical Ensemble. *Mol. Phys.* **1984**, *52* (2), 255-268. DOI: 10.1080/00268978400101201.

(5) Hoover, W. G. Canonical Dynamics: Equilibrium Phase-Space Distributions. *Phys. Rev. A Gen. Phys.* **1985**, *31* (3), 1695-1697. DOI: 10.1103/physreva.31.1695.

(6) Martyna, G. J.; Tuckerman, M. E.; Tobias, D. J.; Klein, M. L. Explicit Reversible Integrators for Extended Systems Dynamics. *Mol. Phys.* **1996**, *87* (5), 1117-1157. DOI: 10.1080/00268979600100761.

(7) Bowers, K. J. C., E.; Xu, H.; Dror, R. O.; Eastwood, M. P.; Gregersen, B. A.; Klepeis, J. L.; Kolossvary, I.; Moraes, M. A.; Sacerdoti, F. D.; Salmon, J. K.; Shan, Y.; Shaw, D. E. Scalable Algorithms for Molecular Dynamics Simulations on Commodity Clusters. *Proceedings of the ACM/IEEE Conference on Supercomputing (SC06)* **2006**, 43. DOI: 10.1109/SC.2006.54.

(8) Abdel-Rahman, S. A.; Talagayev, V.; Pach, S.; Wolber, G.; Gabr, M. T. Discovery of Small-Molecule TIM-3 Inhibitors for Acute Myeloid Leukemia Using Pharmacophore-Based Virtual Screening. *J. Med. Chem.* **2023**, *66* (16), 11464-11475. DOI: 10.1021/acs.jmedchem.3c00960.

(9) Calvo-Barreiro, L.; Talagayev, V.; Pach, S.; Abdel-Rahman, S. A.; Wolber, G.; Gabr, M. T. Discovery of ICOS-Targeted Small Molecules Using Pharmacophore-Based Screening. *ChemMedChem* **2023**, *18* (23), e202300305. DOI: 10.1002/cmdc.202300305.

(10) Schaller, D.; Pach, S.; Wolber, G. PyRod: Tracing Water Molecules in Molecular Dynamics Simulations. *J. Chem. Inf. Model.* **2019**, *59* (6), 2818-2829. DOI: 10.1021/acs.jcim.9b00281.

(11) Bock, A.; Bermudez, M.; Krebs, F.; Matera, C.; Chirinda, B.; Sydow, D.; Dallanoce, C.; Holzgrabe, U.; De Amici, M.; Lohse, M. J.; et al. Ligand Binding Ensembles Determine Graded Agonist Efficacies at a G Protein-coupled Receptor. *J Biol Chem* **2016**, *291* (31), 16375-16389. DOI: 10.1074/jbc.M116.735431.

(12) Wolber, G.; Langer, T. LigandScout: 3-D Pharmacophores Derived from Protein-Bound Ligands and Their Use as Virtual Screening Filters. *J. Chem. Inf. Model.* **2005**, *45* (1), 160-169. DOI: 10.1021/ci049885e.

(13) Wolber, G.; Dornhofer, A. A.; Langer, T. Efficient Overlay of Small Organic Molecules Using 3D Pharmacophores. *J. Comput. Aided Mol. Des.* **2006**, *20* (12), 773-788. DOI: 10.1007/s10822-006-9078-7.

(14) RDKit. RDKit: Open-Source Cheminformatics. *Zenodo 10.5281/zenodo.3732262*. DOI: 10.5281/zenodo.3732262.

(15) Jones, G.; Willett, P.; Glen, R. C.; Leach, A. R.; Taylor, R. Development and Validation of a Genetic Algorithm for Flexible Docking. *J. Mol. Biol.* **1997**, *267* (3), 727-748. DOI: 10.1006/jmbi.1996.0897.

(16) Korb, O.; Stutzle, T.; Exner, T. E. Empirical Scoring Functions for Advanced Protein-Ligand Docking with PLANTS. *J. Chem. Inf. Model.* **2009**, *49* (1), 84-96. DOI: 10.1021/ci800298z.

(17) Fischer, A.; Smiesko, M.; Sellner, M.; Lill, M. A. Decision Making in Structure-Based Drug Discovery: Visual Inspection of Docking Results. *J. Med. Chem.* **2021**, *64* (5), 2489-2500. DOI: 10.1021/acs.jmedchem.0c02227.

(18) Zhao, H. The Science and Art of Structure-Based Virtual Screening. *ACS Med. Chem. Lett.* **2024**, *15* (4), 436-440. DOI: 10.1021/acsmedchemlett.4c00093.

(19) Calvo-Barreiro, L.; Talagayev, V.; Pach, S.; Abdel-Rahman, S. A.; Wolber, G.; Gabr, M. T. Discovery of ICOS-Targeted Small Molecules Using Pharmacophore-Based Screening. *ChemMedChem* **2023**, *18* (23), e202300305. DOI: 10.1002/cmdc.202300305.

(20) Jafari, R.; Almqvist, H.; Axelsson, H.; Ignatushchenko, M.; Lundback, T.; Nordlund, P.; Martinez Molina, D. The cellular thermal shift assay for evaluating drug target interactions in cells. *Nat Protoc* **2014**, *9* (9), 2100-2122. DOI: 10.1038/nprot.2014.138 From NLM Medline.

(21) Kawatkar, A.; Schefter, M.; Hermansson, N. O.; Snijder, A.; Dekker, N.; Brown, D. G.; Lundback, T.; Zhang, A. X.; Castaldi, M. P. CETSA beyond Soluble Targets: a Broad Application to Multipass Transmembrane Proteins. *ACS Chem Biol* **2019**, *14* (9), 1913-1920. DOI: 10.1021/acschembio.9b00399 From NLM Medline.
